# Synthetic lethality of *Mycobacterium tuberculosis* NADH dehydrogenases is due to impaired NADH oxidation

**DOI:** 10.1101/2023.04.10.536268

**Authors:** Yuanyuan Xu, Sabine Ehrt, Dirk Schnappinger, Tiago Beites

## Abstract

Type 2 NADH dehydrogenase (Ndh-2) is an oxidative phosphorylation enzyme discussed as a promising drug target in different pathogens, including *Plasmodium falciparum* and *Mycobacterium tuberculosis* (*Mtb*). To kill *Mtb*, Ndh-2 needs to be inactivated together with the alternative enzyme type 1 NADH dehydrogenase (Ndh-1), but the mechanism of this synthetic lethality remained unknown. Here, we provide insights into the biology of NADH dehydrogenases and a mechanistic explanation for Ndh-1 and Ndh-2 synthetic lethality in *Mtb*. NADH dehydrogenases have two main functions: maintaining an appropriate NADH/NAD+ ratio by converting NADH into NAD+ and providing electrons to the respiratory chain. Heterologous expression of a water forming NADH oxidase (Nox), which catalyzes the oxidation of NADH, allows to distinguish between these two functions and show that Nox rescues Mtb from Ndh-1/Ndh-2 synthetic lethality, indicating that NADH oxidation is the essential function of NADH dehydrogenases for *Mtb* viability. Quantification of intracellular levels of NADH, NAD, ATP, and oxygen consumption revealed that preventing NADH oxidation by Ndh-2 depletes NAD(H) and inhibits respiration. Finally, we show that Ndh-1/ Ndh-2 synthetic lethality can be achieved through chemical inhibition.

**IMPORTANCE:** In 2022, it is estimated that 10.6 million people fell ill, and 1.6 million people died from Tuberculosis (TB). Available treatment is lengthy and requires a multi-drug regimen, which calls for new strategies to cure *Mycobacterium tuberculosis* (*Mtb*) infections more efficiently. We have previously shown that simultaneous inactivation of type 1 (Ndh-1) and type 2 (Ndh-2) NADH dehydrogenase kills *Mtb*. NADH dehydrogenases play two main physiological roles: NADH oxidation and electron entry to the respiratory chain. Here, we show that this bactericidal effect is a consequence of impaired NADH oxidation. Importantly, we demonstrate that Ndh-1/Ndh-2 synthetic lethality can be achieved through simultaneous chemical inhibition, which could be exploited by TB drug development programs.

## INTRODUCTION

Oxidative phosphorylation is an essential cell process for *Mycobacterium tuberculosis* (*Mtb*) – the etiological agent of tuberculosis (TB) - both in replicating and non-replicating conditions (1). This has prompted efforts to identify small molecules that effectively inhibit oxidative phosphorylation, leading to the discovery of the ATP synthase inhibitor bedaquiline (2) - a drug that has contributed to treatment shortening of drug-resistant TB (3).

Given the success of bedaquiline, TB drug development programs have been exploring other possible drug targets in *Mtb*’s oxidative phosphorylation. In this context, type 2 NADH dehydrogenase (Ndh-2) has been discussed as a promising target. Indeed, Ndh-2 inhibition has been proposed as an effective strategy to eradicate infections with other pathogens, like *Plasmodium falciparum* (4) (malaria etiological agent) or *Leishmania sp* (5) (leishmaniases etiological agent). *Mtb’s* genome harbors two genes encoding Ndh-2 enzymes, *ndh* and *ndhA*. Transposon mutant screenings identified *ndh* as required for optimal growth of *Mtb* growth in vitro (6, 7). Moreover, phenothiazines, which are active against replicating and non-replicating bacilli (1), were shown to inhibit Ndh-2 activity and to inhibit respiration (8). These data were thus consistent with an essential role for Ndh-2 in *Mtb’s* oxidative phosphorylation, which together with the absence of a homologue enzyme in humans motivated the identification of specific Ndh-2 inhibitors.

In addition to phenothiazines, which we later showed to inhibit *Mtb’s* in vitro growth independently of Ndh-2 (9), multiple small molecules have been proposed to specifically inhibit Ndh-2 in *Mtb:* quinolones (10), quinolinyl pyrimidines (11), molecules with thioquinazoline and tetrahydroindazole cores (12), diphenyleneiodonium analogues (13), 2-Mercapto-Quinazolinones (14), 7-phenyl benzoxaborole compound series (15), and, more recently, tricyclic spirolactams (16). However, the activity of these compounds against *Mtb* was only demonstrated in in vitro conditions and an Mtb strain in which both genes encoding Ndh-2 enzymes have been deleted (*Mtb Δndh-2*) is only mildly attenuated in a mouse model of infection, suggesting that inhibition of Ndh-2 alone will not kill *Mtb* during infection (9).

In addition to *ndh* and *ndhA*, the *Mtb* genome also contains the *nuo* operon, which encodes a nonessential type 1 NADH dehydrogenase (Ndh-1) (1). Unsurprisingly, Ndh-1 inhibitors do not restrict *Mtb* growth in vitro, even at high concentrations (1, 9). Nevertheless, Ndh-1 can compensate for the absence of Ndh-2 activity and support *Mtb*’s growth in vitro and in a mouse model of infection (9). *Mtb* lacking both the *nuo* operon and *ndh* is attenuated in vivo, suggesting that a mutant devoid of NADH dehydrogenase activity might not be viable (17). Consistent with this, the Ndh-1 inhibitor rotenone killed Mtb Δ*ndh-2* in vitro, thus confirming Ndh-1 and Ndh-2 synthetic lethality (9). Here, we show that this synthetic lethality is due to an impaired NADH oxidation. Moreover, we demonstrate that Ndh-1 and Ndh-2 synthetic lethality can be achieved through chemical inhibition.

## RESULTS

### NADH dehydrogenases synthetic lethality is rescued by the heterologous expression of a NADH oxidase

We have previously demonstrated that the chemical inhibition of Ndh-1 kills Mtb Δ*ndh-2* (9). In this work, we sought to understand the molecular mechanism of this synthetic lethality. NADH dehydrogenases play an important role in oxidative phosphorylation and in NADH oxidation into NAD^+^. Previously, we showed that the heterologous expression of Nox, an enzyme that uses oxygen to to convert NADH into NAD^+^ and water (Fig. 1A), rescued the sensitivity of Mtb Δ*ndh-2* to highly reduced carbon sources (9). We thus hypothesized that Nox could deconvolute the importance of NADH dehydrogenases in oxidative phosphorylation versus NADH oxidation. To test this, we determined the minimal inhibitory concentration (MIC) and minimal bactericidal concentration (MBC) of three different Ndh-1 inhibitors, pyridaben, piericidin A and rotenone. As previously observed, wild-type and complemented strain (Δ*ndh-2*::comp) were resistant to Ndh-1 inhibitors, while Δ*ndh-2* was hypersusceptible to all Ndh-1 inhibitors (9) (Fig. 1B). Interestingly, expression of Nox in Δ*ndh-2* (Δ*ndh-2::nox*; Fig. S1) conferred increased resistance to all Ndh-1 inhibitors (Fig. 1B). This effect was not observed with isoniazid (INH) or ethambutol, showing that Nox expression does not confer unspecific resistance to antibiotics (Fig. 1C). Next, we measured bacterial viability upon treatment with the same panel of Ndh-1 inhibitors. As expected, all Ndh-1 inhibitors killed Δ*ndh-2*, but not the wild-type and Δ*ndh-2*::comp (Fig. 2A). Expression of Nox rescued Mtb Δ*ndh-2* from killing by rotenone and, to a lesser degree, pyridaben and piericidn A (Fig. 2A). The variability observed among different Ndh-1 inhibitors might be due to a saturation of Nox activity to overcome Ndh-1 inhibition and/or compound unspecific effects (Fig. 2A). This rescue effect was not observed in the control compounds INH and ethambutol (Fig. 2B).

**FIG 1.**
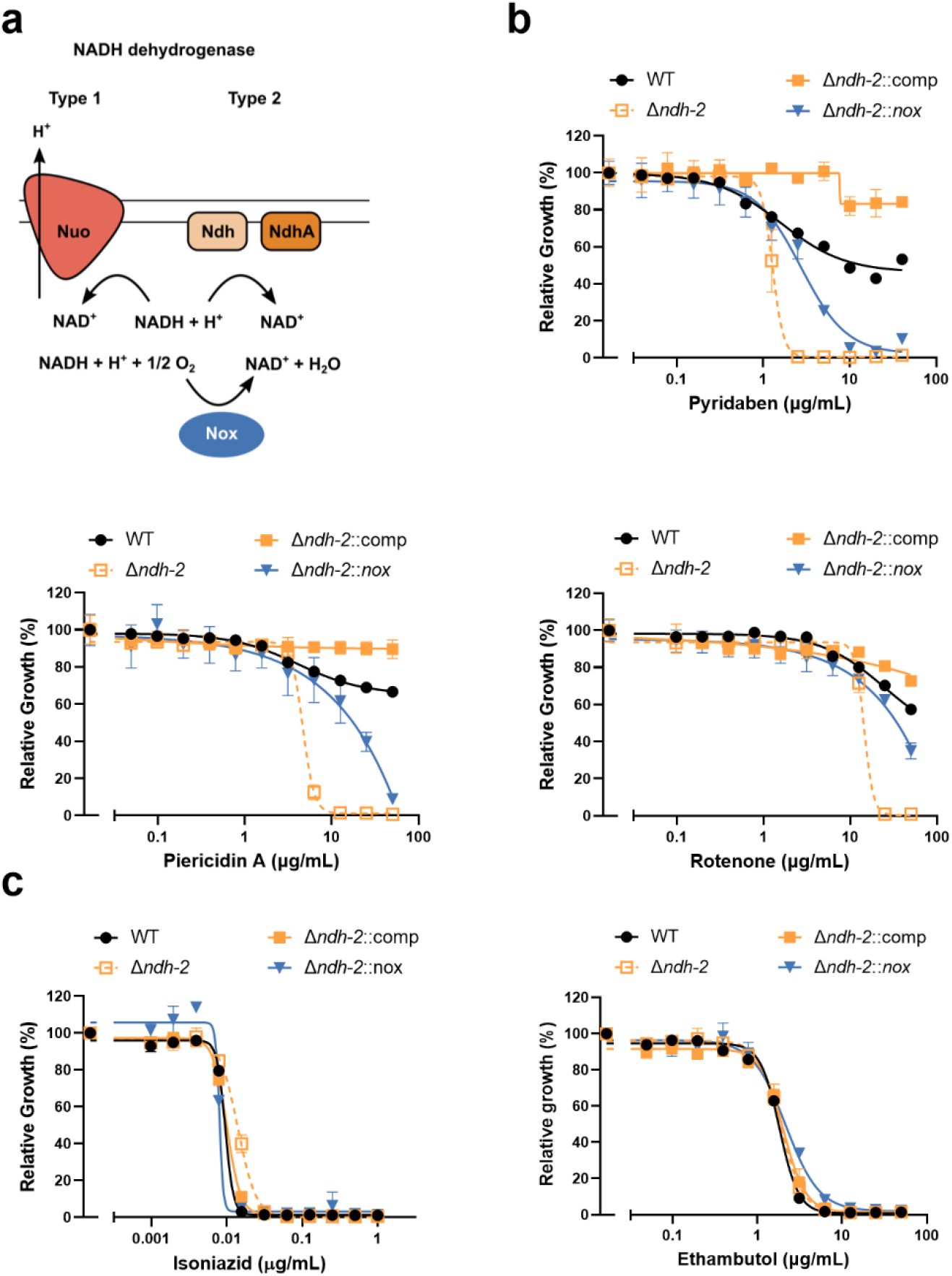
Minimal inhibitory concentration measurements to test if expression of the water-forming NADH oxidase Nox (a) confers resistance to Ndh-1 inhibition in a Δ*ndh-2* genetic background. Wild-type (WT), Δ*ndh-2*, complemented strain (Δ*ndh-2*::comp) and Δ*ndh-2* expressing *nox* (Δ*ndh-2::nox*) were grown in modified Sauton’s medium and tested for susceptibility to the Ndh-1 inhibitors pyridaben, piericidin A and rotenone (b) and to isoniazid and ethambutol as controls (c). Results correspond to OD_580nm_ normalized to no-drug control at day 14 of treatment. Data are averages of technical triplicates. Error bars correspond to standard deviation. These data are representative of 3 independent experiments.

**FIG 2.**
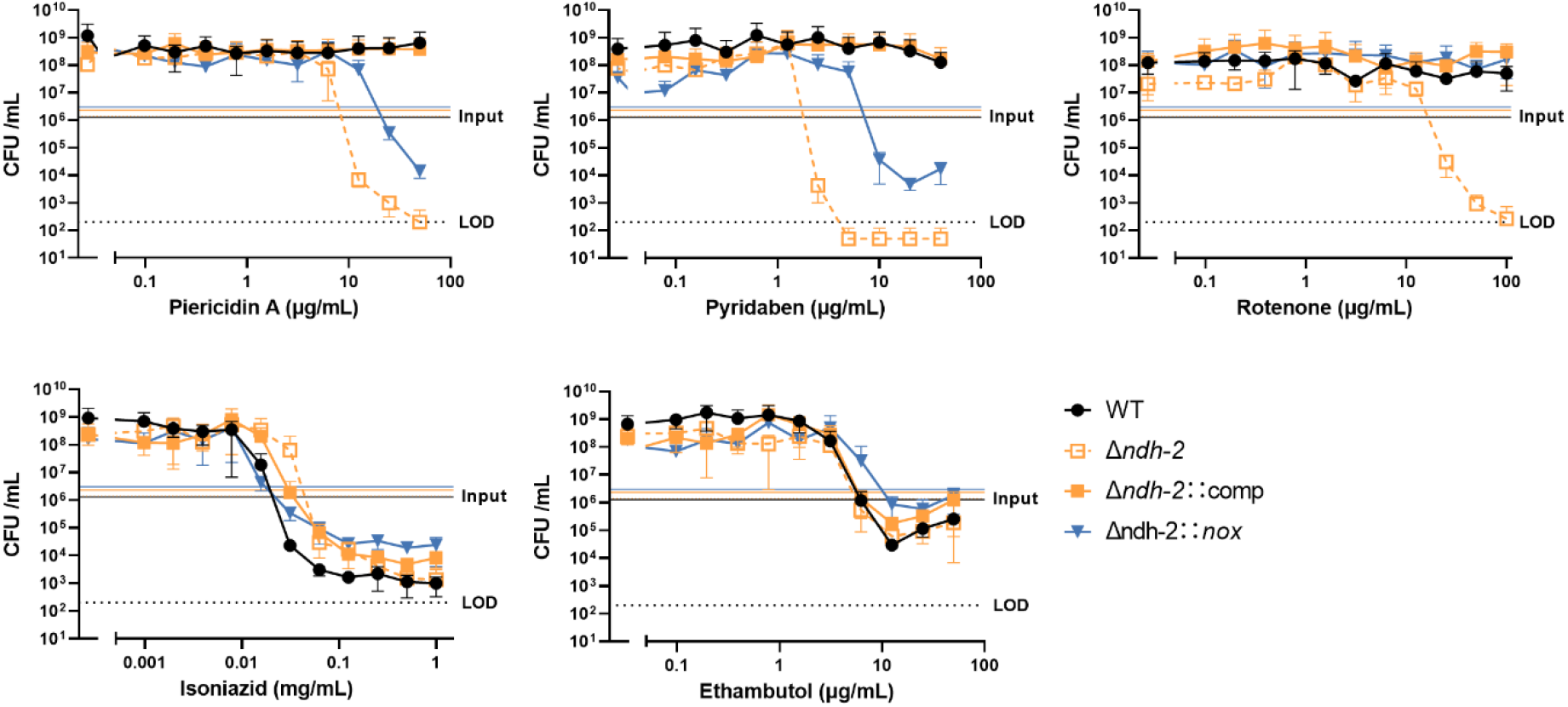
Minimal bactericidal activity measurements. Wild-type (WT), Δ*ndh-2*, complemented strain (Δ*ndh-2*::comp) and Δ*ndh-2* expressing *nox* (Δ*ndh-2::nox*) were grown in modified Sauton’s medium and treated with Ndh-1 inhibitors (pyridaben, piericidin A and rotenone), isoniazid and ethambutol for 14 days. Results correspond to colony forming units (CFU). Data are averages of technical triplicates. Error bars correspond to standard deviation. This experiment is representative of 3 independent experiments. LOD = limit of detection.

These data show that an alternative mechanism of NADH oxidation, provided by Nox, is capable of rescuing *Mtb* viability from a deficient NADH dehydrogenase activity. This observation indicates that the *Mtb* respiratory chain can compensate for the lack of functional NADH dehydrogenases, but the bacilli do not have an effective alternative mechanism for NADH oxidation.

### Inhibition of Ndh-1 in Mtb Δ*ndh-2* leads to depletion of intracellular NAD(H) pools

One of the possible consequences for an impairment in NADH oxidation is a disruption of the cellular redox balance in the form of the NADH/NAD+ ratio. To test this hypothesis, we quantified intracellular NAD(H) pools in response to treatment with pyridaben (Pyr) at 2 μg/ml and 4 μg/ml. We collected samples after 24h treatment to observe the primary effects of Ndh-1 inhibition in Δ*ndh-2* (growth arrest was only observed 48 h post-treatment; Fig. S2). As previously observed, Δ*ndh-2* displayed a higher NADH/NAD+ ratio than the wild-type and Δ*ndh-2*::comp in medium compatible with growth (vehicle control) (9), which indicates that Ndh-2 has a more prominent role than Ndh-1 in redox homeostasis (Fig. 3A). However, Ndh-1 inhibition had a marginal effect on the NADH/NAD+ ratio of Mtb Δ*ndh-2* (1.4x increase with the highest Pyr concentration). Wild-type, Δ*ndh-2*::comp and Δ*ndh-2::nox* did not show significant alterations in NADH/NAD+ ratio upon treatment with Pyr. This suggests that a disruption of NADH/NAD+ ratio is not in the basis of Ndh-1/ Ndh-2 synthetic lethality. Next, we analyzed the individual intracellular concentrations of NADH and NAD+. Consistent with a defect in NADH oxidation, Pyr treatment depleted NAD+ (NADH dehydrogenases product) intracellular levels in Δ*ndh-2* in a dose-dependent manner (Fig. 3B). Curiously, Pyr treatment slightly lowered Δ*ndh-2* intracellular NADH (NADH dehydrogenases substrate) levels. Wild-type, Δ*ndh-2*::comp and Δ*ndh-2::nox* intracellular NAD(H) levels did not show a Pyr dose dependent effect. This can explain the relatively stable NADH/NAD+ ratio in Δ*ndh-2* and shows that a depletion of total NAD(H) intracellular pools is an effect of Ndh-1/ Ndh-2 synthetic lethality.

**FIG 3.**
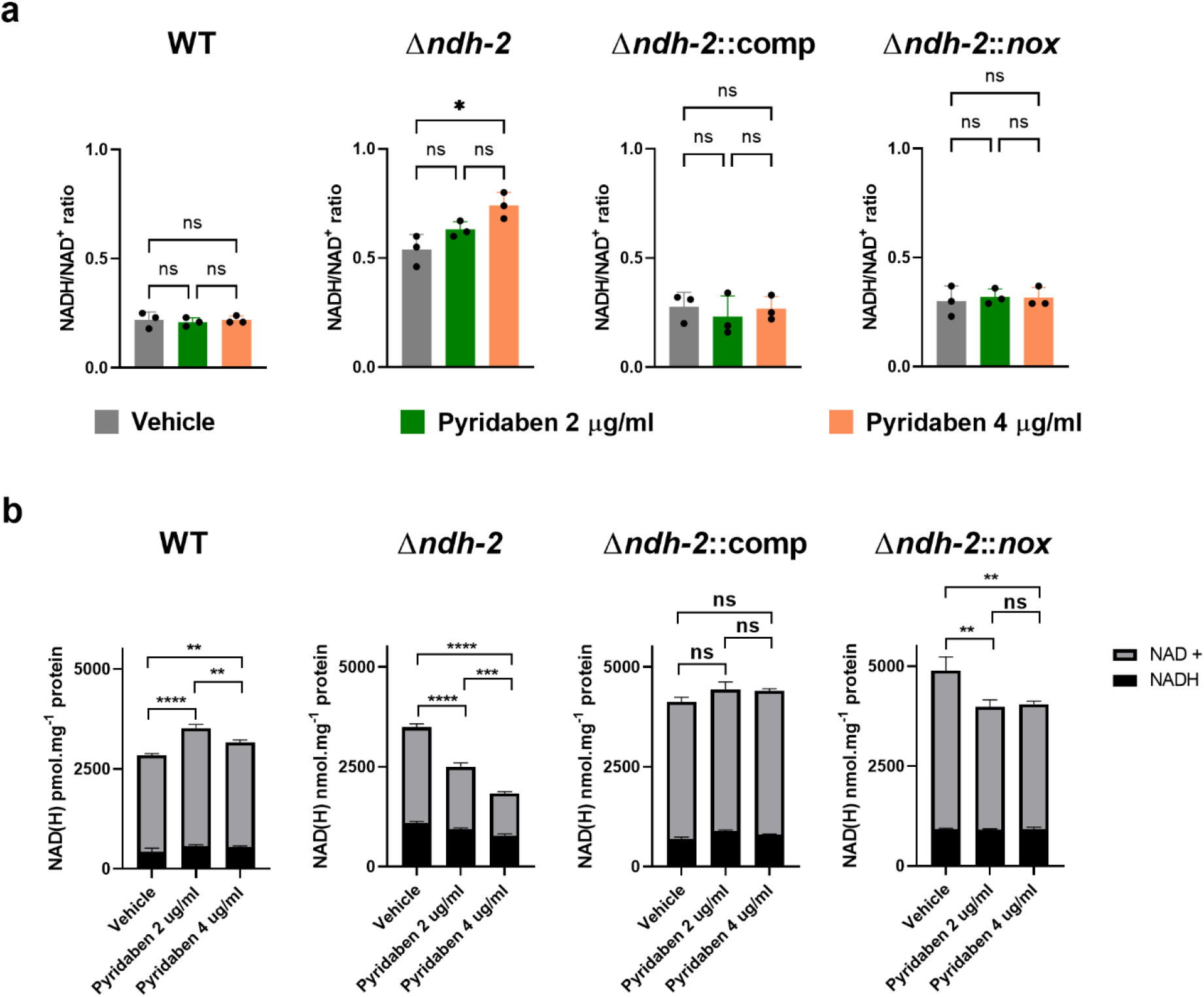
Intracellular NAD(H) concentrations. Wild-type (WT), Δ*ndh-2*, complemented strain (Δ*ndh-2*::comp) and Δ*ndh-2* expressing *nox* (Δ*ndh-2::nox*) were grown in modified Sauton’s medium until mid-exponential phase (OD_580nm_ of 0.5) and challenged with 2 concentrations of pyridaben (2 μg/ml and 4 μg/ml), or the vehicle (DMSO) for 24 hours. (a) NADH/NAD+ ratio measurements are averages of 3 independent replicates. Error bars correspond to standard deviation. (b) NADH and NAD+ intracellular concentrations. Data are averages of 3 technical replicates. Error bars correspond to standard deviation. These data are representative of 3 independent experiments. Statistical significance was assessed by one-way ANOVA followed by post hoc test (Tukey test; GraphPad Prism). \*\**P* ≤ 0.01; \*\*\**P*≤ 0.001; \*\*\*\**P*≤ 0.0001 ns - not significant.

Our data show that an impaired NADH dehydrogenase activity results in reduced intracellular NAD(H) pools, which has been shown to exert a bactericidal effect during infection (18).

### Defective NADH dehydrogenase activity impacts oxidative phosphorylation indirectly

To assess if a defective overall NADH dehydrogenase activity compromises *Mtb* respiration, we measured the oxygen consumption rate (OCR) in bacterial suspensions (PBS tyloxapol 0.05%) treated with Pyr (2 μg/ml and 4 μg/ml). In Mtb Δ*ndh-2*, Pyr treatment resulted in a concentration dependent inhibition of oxygen consumption (Fig. 4A): ~15% inhibition with Pyr 2 μg/ml and ~48% inhibition with Pyr 4 μg/ml. Wild type and Δ*ndh-2*::comp OCR was insensitive to Pyr treatment. These results are compatible with a prominent role of NADH dehydrogenases in the input of electrons to the respiratory chain. However, Δ*ndh-2::nox* was also insensitive to Pyr treatment, indicating that the impact on respiration is indirect and stems from an impaired NADH oxidation.

**FIG 4.**
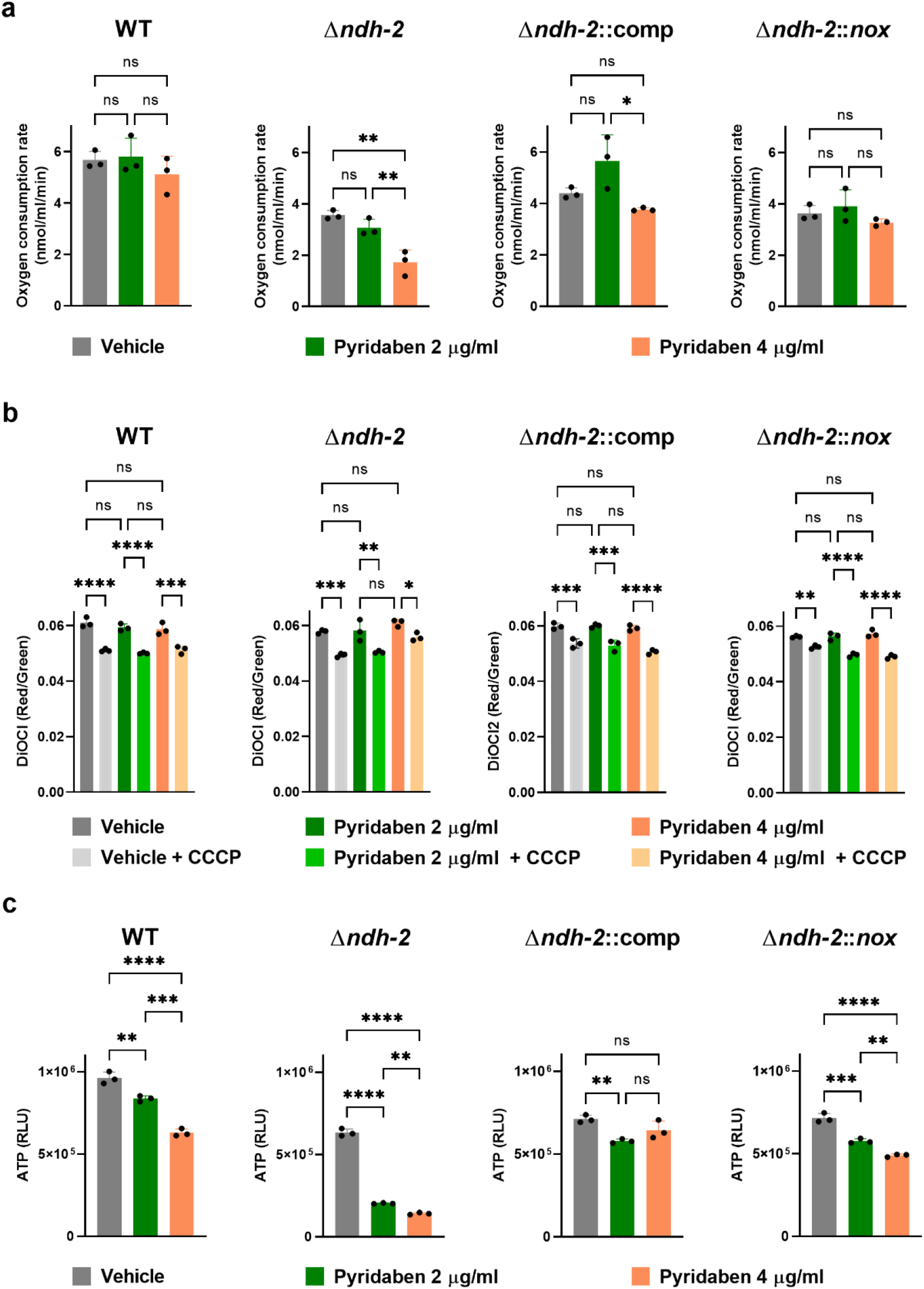
Quantification of oxidative phosphorylation related variables. (a) Oxygen consumption rate measurements. Glycerol (carbon source) and pyridaben (2 μg/ml or 4 μg/ml) or vehicle (DMSO) were added to bacteria suspensions in PBS-tyloxapol (OD_580nm_ of 0.5) until a stable oxygen consumption rate was achieved. (b) Membrane potential determinations using the fluorescent probe DiOC_2_. Strains were cultured in modified Sauton’s medium and treated with pyridaben (2 μg/ml or 4 μg/ml) or vehicle (DMSO) for 24 hours. Protonophore carbonyl-cyanide 3-chlorophenylhydrazone (CCCP) was used as a control for membrane depolarization. Results are averages of technical replicates. Error bars correspond to standard deviation. Data are representative of 3 independent experiments. (c) Intracellular ATP levels. Strains were cultured in modified Sauton’s medium and treated with pyridaben (2 μg/ml or 4 μg/ml) or vehicle (DMSO) for 24 hours. Results are averages of technical replicates. Error bars correspond to standard deviation. Data are representative of 3 independent experiments. Statistical significance was assessed by one-way ANOVA followed by post hoc test (Tukey test; GraphPad Prism). \**P* ≤ 0.05; \*\**P* ≤ 0.01; \*\*\**P* ≤ 0.001; \*\*\*\**P* ≤ 0.0001 ns - not significant.

Next, we measured membrane potential (Fig. 4B) and intracellular ATP levels (Fig. 4C) after a 24 h treatment with Pyr (2 μg/ml and 4 μg/ml), using the same experimental design previously described (Fig. S2). This revealed that Pyr treatment did not depolarize membrane potential of Mtb Δ*ndh-2*. Nevertheless, Pyr lowered intracellular ATP levels in Δ*ndh-2*, which is consistent with a partial inhibition of the respiratory chain. Wild-type, Δ*ndh-2*::comp, and Δ*ndh-2::nox* showed only a modest, but statistically significant effect, on ATP intracellular levels. Of note, the OCR and intracellular ATP levels of Mtb Δ*ndh-2* were lower than those of wild-type in the vehicle control situation, which again argues in favor of Ndh-2 taking a more prominent role in *Mtb’s* physiology. Also, Δ*ndh-2*::comp did not show OCR and intracellular ATP levels similar to the wild-type, which may be because this strain is only expressing *ndh*.

In summary, a defective overall NADH dehydrogenase activity negatively impacts *Mtb’s* oxidative phosphorylation. However, this stems from an impaired NADH oxidation, which may inhibit multiple pathways necessary and ultimately impacting respiration.

### Simultaneous chemical inhibition of Ndh-1 and Ndh-2 kills *Mycobacterium tuberculosis*

With cytochrome bc1-aa3 oxidase inhibitor Q203 (Telacebec) (19) in clinical trials (20) and the discovery of a potent cytochrome bd oxidase inhibitor (21), synthetic lethality of *Mtb’s* terminal oxidases has been proposed as a new strategy to treat infections (22). We sought to verify if the same strategy could be applied to NADH dehydrogenases. To show that a bactericidal effect could be obtained through simultaneous chemical inhibition of both enzymes, we determined the kill kinetics of wild-type *Mtb* following treatment with the Ndh-1 inhibitor Pyr and the Ndh-2 inhibitor 2-Mercapto-Quinazolinone (2-MQ) DDD00853663 (14) individually or in combination (Fig. 5). Drug concentrations were based on MIC values for these compounds, which we determined in a previous study (9). To further support that a defective NADH oxidation is the cause of the synthetic lethality, we exposed wild-type *Mtb* expressing *nox* (Fig. S1) to the same conditions. INH was chosen as a control compound. As expected, in these experimental conditions, wild-type is not susceptible to Ndh-1 or Ndh-2 inhibition alone (Ndh-2 is only essential when long-chain fatty acids are present in the medium(9)). However, simultaneous chemical inhibition of Ndh-1 and Ndh-2 led to a bactericidal effect. Consistent with our proposed mechanism, Nox expression rescued *Mtb’s* viability upon simultaneous treatment with Ndh-1 and Ndh-2 inhibitors. Importantly, this rescue effect was not observed with INH.

**FIG 5.**
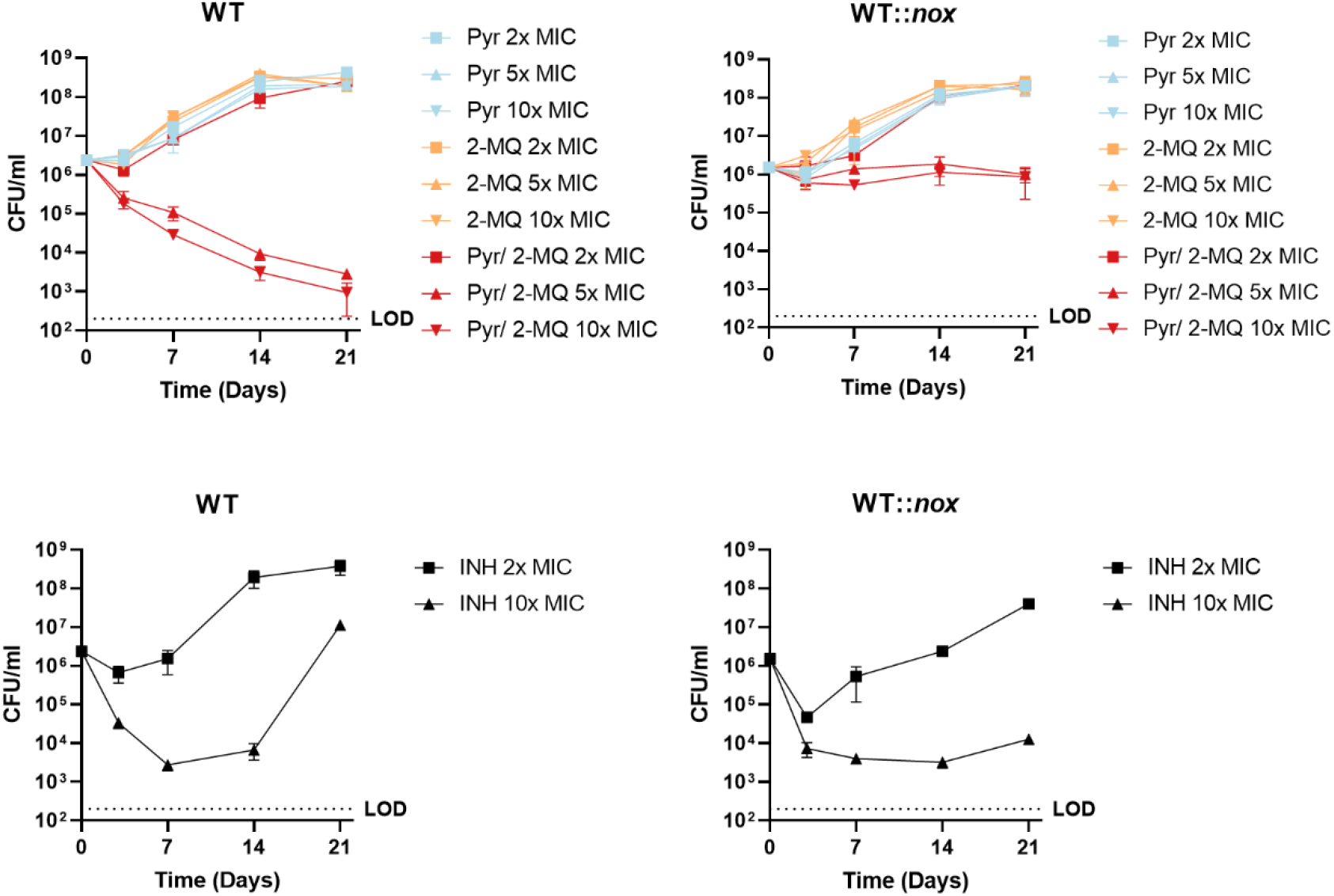
Kill curves. Wild type (WT) and wild type expressing *nox* (WT::*nox*) were treated with pyridaben (pyr) or the 2-Mercapto-Quinazolinone (2-MQ) DDD00853663 alone or in combination at concentrations (2x, 5x or 10x MIC) based on Pyr MIC in Δ*ndh-2*, or 2-MQ MIC in medium with oleic acid. Isoniazid (INH) was used as a control drug at 2x and 10x MIC. Results are averages of technical triplicates. Error bars correspond to standard deviation. These data are representative of 3 independent experiments. LOD = limit of detection.

Our results show that it is possible to exploit the synthetical lethality of *Mtb’s* NADH dehydrogenases through chemical inhibition and confirmed that the bactericidal effect is due to impaired NADH oxidation.

## DISCUSSION

The *Mtb* respiratory chain is a highly branched and plastic cellular process composed of nine respiratory dehydrogenases and four terminal oxidoreductases. The respiratory enzyme Ndh-2 was deemed to be required for in vitro growth (6, 7) and a major electron entry point to the respiratory chain (8). We have previously shown that Ndh-2 is conditionally essential for in vitro growth depending on the presence of highly reduced carbon sources and that Ndh-1 can compensate for the lack of Ndh-2 during mouse infection (9). Here, we sought to understand the molecular mechanism of Ndh-2/ Ndh-1 synthetic lethality.

NADH dehydrogenases have two major functions: 1) NADH oxidation and 2) input of electrons to the respiratory chain. We have previously shown that a water forming NADH oxidase (Nox) is functional in *Mtb* (9) and reasoned that this enzymatic activity could disambiguate the roles of NADH dehydrogenases in NADH oxidation and respiration. We demonstrate that Nox expression rescues Mtb Δ*ndh-2* (lacking *ndh* and *ndhA*) from the bactericidal effect of Ndh-1 chemical inhibition. NADH dehydrogenase activity is, thus, essential for bacilli viability, at least in the tested conditions, due to the role in NADH oxidation rather than being the major donor of electrons to the respiratory chain.

The measurement of intracellular NAD(H) pools in Δ*ndh-2* challenged with chemical Ndh-1 inhibitors showed a depletion of NAD+, which is consistent with an impaired NADH oxidation. However, depletion of NAD+ was not accompanied by the accumulation of NADH. On the contrary, intracellular NADH levels decreased in response to Pyr treatment, resulting in only a modest increase of the NADH/NAD+ ratio, which is in contrast with what is observed in other bacteria (23). Instead, Ndh-1 chemical inhibition in Δ*ndh-2* led to a marked decrease in the intracellular NAD(H) pools, an effect that was shown to be bactericidal to *Mtb* in vitro and during infection (18). Importantly, Nox restored the NAD(H) concentrations of Mtb Δ*ndh-2* to wild-type levels. These phenotypes are thus consistent with a model in which the absence of an efficient NADH oxidation system triggers a mechanism that avoids NADH accumulation to balance NADH/NAD+ ratio; however, in doing so, it traps the bacilli in a feedback loop that leads to NAD(H) intracellular pool depletion. The nature of this mechanism is unknown, but one possibility could be the NADH-inducible nudix hydrolase RenU that cleaves NADH with high specificity (24).

The evaluation of bioenergetic parameters, namely OCR, membrane potential and intracellular ATP levels were consistent with a negative impact of deficient overall NADH dehydrogenase activity in oxidative phosphorylation. However, that these phenotypes were complemented by the expression of *nox*, argues in favor of an indirect effect stemming from an impairment in NADH oxidation. *Mtb* expresses multiple dehydrogenases capable of reducing menaquinone, including succinate dehydrogenases that were recently shown to be essential for growth (25), which may compensate the lack of NADH dehydrogenase activity. This is in contrast with other bacterial pathogens like *Streptococcus agalactiae* – a causal agent of sepsis in a newborns and immunocompromised adults – which uses NADH dehydrogenases as the main entry point of electrons to the respiratory chain (26).

Treatment of *Mtb* with a combination of Ndh-1 and Ndh-2 inhibitors showed that a bactericidal effect can be achieved by chemical inhibition. Importantly, expression of Nox rescued wild-type *Mtb* from Ndh-1/Ndh-2 synthetic lethality, further confirming that NADH oxidation is the main biological function of NADH dehydrogenases regarding bacterial viability. Several potent Ndh-2 inhibitors have been identified (10–16). Δ*ndh-2* susceptibility to long-chain fatty acids in the medium, an important carbon source during infection, and to hypoxic conditions (9) argues for Ndh-2 inhibitors to be effective in specific microenvironments like the necrotic center of caseating granulomas. However, that an *Mtb Δndh-2* grows and survives during mouse infection (9) shows that Ndh-2 inhibition may not be effective in all infection microenvironments. To circumvent this limitation, Ndh-1/Ndh-2 synthetic lethality could be exploited. This is because simultaneous inhibition of both NADH dehydrogenases leads to a compound effect of at least two conditions that can contribute to a bactericidal effect: intracellular NAD(H) pool depletion (18) and lower intracellular ATP levels (1). The main challenge to this approach would be the identification of inhibitors with high affinity for *Mtb’s* Ndh-1 and low affinity to the homologue enzyme in humans (complex I). The currently known Ndh-1 inhibitors interact with the quinone-binding pocket (27) of species across the tree of life, including humans, making them toxic. However, one could make a case for such an endeavor given the successful identification of inhibitors to *Mtb* enzymes that have homologues in humans, like bedaquiline (2) (ATP synthase) and Q203 (cytochrome *bc1* complex) (19). Moreover, the structure of the mycobacterial Ndh-1 enzyme complex was recently solved (28), which could help in of the design of Ndh-1 inhibitors.

Our work unveiled NADH oxidation as the essential function of NADH dehydrogenases for maintaining *Mtb* viability. Deficient overall NADH dehydrogenase activity was associated with depletion of NAD(H) intracellular tools and inhibition of oxidative phosphorylation. Finally, we showed that Ndh-1/Ndh-2 synthetic lethality can be exploited to kill *Mtb* by chemically inactivating these enzymes.

## METHODS

### Growth conditions and strains

*Mtb* strains (H37Rv genetic background) were cultured in modified Sauton’s minimal medium: 0.05% (w/v) potassium dihydrogen phosphate, 0.05% (w/v) magnesium sulfate heptahydrate, 0.2% (w/v) citric acid, 0.005% (w/v) ferric ammonium citrate, 0.0001% (w/v) zinc sulfate and supplemented with 0.05% (v/v) tyloxapol, 0.4% (w/v) glucose, 0.2% (v/v) glycerol, and ADNaCl with fatty-acid-free BSA (Roche). For modified Sauton’s solid medium, 1.5% (w/v) bactoagar (BD) was added; also, glycerol was added at 0.5% (w/v). Solid medium was used for transformations and colony forming units (CFU) outgrowth. When necessary, antibiotics were added to cultures (final concentrations): hygromycin 50 μg/ml, kanamycin 25 μg/ml, streptomycin 50 μg/ml, and zeocin 12.5 μg/ml. Δ*ndh-2*, Δ*ndh-2* complemented (expression of a native copy of *ndh*) and Δ*ndh-2::nox* (strain with an integrative plasmid expressing an *Mtb* codon adapted version of *nox* from *Lactococcus lactis* under the transcriptional control of the promoter Ptb38 - pGMCgS-0×-Ptb38-NOX-FLAG-SD1) were generated in a previous study(9). For this study, we have transformed wild-type with the integrative plasmid pGMCgS-0×-Ptb38-NOX-FLAG-SD1 to generate *WT::nox*.

### Immunodetection

WT, Δ*ndh-2*, and Δ*ndh-2::nox* and WT::*nox* cultures were grown until mid-exponential phase. Bacteria were washed with PBS with 0.05% tyloxapol and resuspended in 500 μL PBS with 1xprotease inhibitor cocktail (Roche). Bacterial lysis was performed by bead-beating three times at 4500 rpm for 60s with 0.1 mm Zirconia/Silica beads. Supernatants were then filtered through a 0.2-μm SpinX column (Corning). Protein concentrations were determined using Qubit Protein Assay Kit (Invitrogen). We used 30 μg protein, separated the protein extracts through SDS-PAGE, transferred to nitrocellulose membranes and proceeded with the incubation with primary antibody, anti-Flag (Sigma-Aldrich, at 1:1000 dilution) and anti-PrcB (Sigma-Aldrich, at 1:1000 dilution) at room temperature for 2 hours. The secondary antibodies goat anti-mouse IgG, (Thermo Fisher, DyLight 800), and Donkey anti-Rabbit IgG (LI-COR Biosciences, IRDye 680LT) were used at a 1:10000 dilution and incubated at room temperature for 30 min. Immunodetection was performed in an Odyssey Infrared Imaging System (LI-COR Biosciences).

### Minimal inhibitory/bactericidal concentration and kill curves

For minimal inhibitory concentration (MIC) assays, strains were grown in modified Sauton’s medium until mid-exponential phase (OD_580nm_ of 1) and resuspended in fresh medium as single bacteria suspensions. 96-well plates with 11 concentration drugs (2-fold dilutions) plus a no-drug control in triplicates (dispensed by D300e Digital Dispenser - HP) were seeded with 200 μl of single bacteria suspensions. DMSO was normalized across wells to 1% (v/v). Plates were incubated for 14 days and OD_580nm_ was then recorded. Results were presented as a percentage of drug over no-drug control. To estimate minimal bactericidal concentration (MBC), we took samples from the MIC plates and serially diluted the culture in PBS tyloxapol 05% (v/v). Modified Sauton’s solid medium was used for outgrowth. Kill curves were performed in 96-well plates following the same culture conditions described for MIC/MBC. Drugs were dispensed by a D300e Digital Dispenser (HP), with DMSO being normalized to 1% (v/v) across wells. Samples for CFU determination were harvested at days 3, 7, 14 and 21. Modified Sauton’s solid medium was used for outgrowth.

### NAD(H) quantification, membrane potential and intracellular ATP levels

NAD(H) quantification, membrane potential and ATP intracellular levels were determined using the same experimental design. *Mtb* strains were cultured (10 ml; unvented T-75 flasks; with rotation 100 rpm) in modified Sauton’s minimal medium until mid-exponential phase (OD_580nm_ of 1) and resuspended in fresh medium at an OD_580nm_ of 0.5; DMSO, pyridaben 2 μg/ml and 4 μg/ml were then added to the cultures (DMSO was normalized to the highest concentration across conditions). Bacteria were harvested after a 24h treatment (time point that precedes the inflexion point of Δ*ndh-2* growth treated with pyridaben). NAD(H) intracellular concentration was quantified with the commercial kit Fluoro NAD (Cell Technnology) and following the manufacturer’s instructions. NAD(H) intracellular concentrations were normalized by protein content (Qubit Protein Assay Kit - Invitrogen). For membrane potential, bacteria were resuspended in fresh medium (same experimental conditions) with a final OD_580nm_ of 1 and treated with 15 μM DiOC2 3 for 30 min (room temperature). Protonophore carbonyl-cyanide 3-chlorophenylhydrazone (CCCP) was added at a final concentration of 50 μM to provide a depolarized membrane control. Samples were then washed in fresh media (same experimental conditions) and 200 μl in triplicate were dispensed in a black clear bottom 96-well plates (Costar). Fluorescence was recorded in a SpectraMax M5 spectrofluorimeter (Molecular Devices): green fluorescence (488 nm/530 nm) and red fluorescence (488 nm/610 nm). A shift to red is synonymous of dye aggregation caused by membrane potential. Membrane potential was estimated as a ratio of red fluorescence over green fluorescence. ATP intracellular levels were determined using the commercial kit BacTiter-Glo™ (Promega) and following the manufacturer’s instructions.

### Oxygen consumption rate quantification

*Mtb* strains were cultured in modified Sauton’s until mid-exponential phase (OD_580nm_ of 1) and resuspended in pre-heated (37 °C) PBS tyloxapol 0.5% (v/v) to a final OD of 0.5. A Clark-type electrode system (Oxytherm+; Hansatech) was used to measure dissolved oxygen concentration. After calibrating the instrument following the manufacturer’s instructions, we added the following components to the chamber: 950 μl of bacteria suspension, 25 μl glycerol (1 M) and 25 μl of pyridaben (40x of treatment concentration) or DMSO (vehicle). Oxygen concentration was followed and recorded using the software Oxytrace+.

## ACKNOWLEDGEMENTS

We thank Peter Finin from NIH/NIAID for technical advice and useful discussions. Helena Boshoff for generously providing an aliquot of the compound DDD00853663. This work was supported by the NIH grant P01AI143575.

