## Supplementary Figures for "Synthetic lethality of *Mycobacterium tuberculosis* NADH dehydrogenases is due to impaired NADH oxidation"

#### Supplementary Information

### *Mycobacterium tuberculosis* type 1 and type 2 NADH dehydrogenase synthetic lethality is due to impaired NADH re-oxidation

Yuanyuan Xu, Sabine Ehrt, Dirk Schnappinger, Tiago Beites

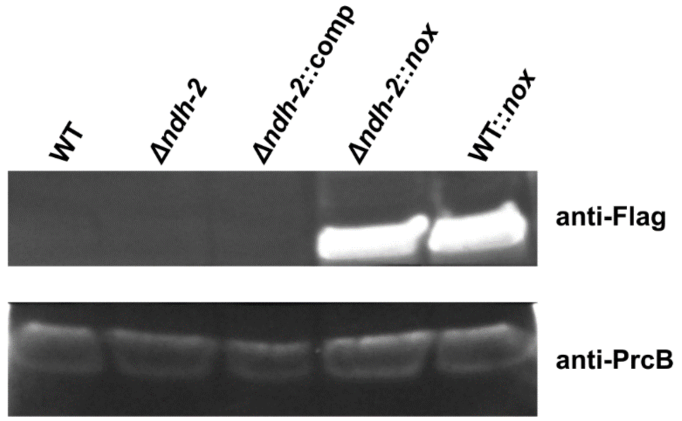


FIG S1 Immunodetection of Nox-Flag expression. Nox-Flag was detected with an anti-flag antibody. PrcB was used as a loading control.

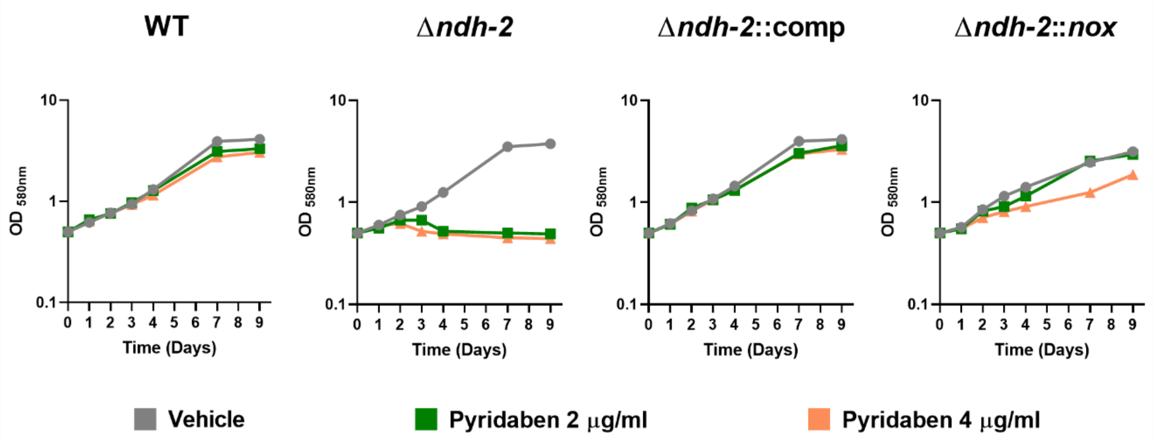


FIG S2 Growth kinetics upon treatment with Pyridaben (2 μg/ml and 4 μg/ml) or vehicle DMSO. These results are representative of 5 independent experiments.
